# Inferring protein fitness landscapes from laboratory evolution experiments

**DOI:** 10.1101/2022.09.01.506224

**Authors:** Sameer D’Costa, Emily C. Hinds, Chase R. Freschlin, Hyebin Song, Philip A. Romero

## Abstract

Directed laboratory evolution applies iterative rounds of mutation and selection to explore the protein fitness landscape and provides rich information regarding the underlying relationships between protein sequence, structure, and function. Laboratory evolution data consist of protein sequences sampled from evolving populations over multiple generations and this data type does not fit into established supervised and unsupervised machine learning approaches. We develop a statistical learning framework that models the evolutionary process and can infer the protein fitness landscape from multiple snapshots along an evolutionary trajectory. We apply our modeling approach to dihydrofolate reductase (DHFR) laboratory evolution data and the resulting landscape parameters capture important aspects of DHFR structure and function. We use the resulting model to understand the structure of the fitness landscape and find numerous examples of epistasis but an overall global peak that is evolutionarily accessible from most starting sequences. Finally, we use the model to perform an *in silico* extrapolation of the DHFR laboratory evolution trajectory and computationally design proteins from future evolutionary rounds.

## Introduction

The mapping from protein sequence to function forms a high-dimensional protein fitness landscape. Knowledge of this landscape is important for understanding and modeling natural evolution, diagnosing genetic diseases, and designing new proteins for applications in biotechnology, human health, and chemistry. This landscape is shaped by highly complex protein conformations, dynamics, and biophysical/biochemical mechanisms, and is defined over an astronomically large number of possible protein sequences. While the sequence-function mapping is challenging to model from a physical perspective, approaches from statistics and machine learning can be leveraged to infer the underlying landscape from sparsely sampled experimental and evolutionary data [1–4].

The statistical approaches to model the protein fitness landscape are built around two common data types that provide either labeled or unlabeled data. Labeled protein data consist of a set of amino acid sequences and how each of those sequences map to a particular protein property of interest, such as thermostability, enzyme activity, or binding affinity. These sequence-function data are commonly generated using protein mutagenesis libraries and medium- or high-throughput assays to assign functional labels [5, 6]. Supervised learning approaches such as linear regression or more sophisticated non-linear models can learn from labeled sequence-function data to infer the mapping from sequence to function [7–10]. Unlabeled protein data consist of natural protein sequences taken from genomic and metagenomic sequencing databases. Unsupervised learning approaches can learn from this unlabeled protein data to infer the fitness landscape [11–13]. Direct coupling analysis (DCA) is an important class of unsupervised learning methods that learn residue coevolution patterns from multiple sequence alignments of related sequences [14, 15]. The DCA approach has been used to predict the three-dimensional structures of proteins [16–18], model the effects of mutations [19], and design new proteins [20].

Directed laboratory evolution applies iterative rounds of mutation and selection to explore the protein fitness landscape [7]. As a population evolves it samples diverse regions of sequence space and generates evolutionary trajectories that can be used to understand the structure of the fitness landscape. Laboratory evolution data consist of protein sequences sampled from evolving populations over multiple sequential generations. These data don’t naturally fit into established supervised or unsupervised learning paradigms. Previous work has treated laboratory evolution data similar to natural evolution data and performed unsupervised DCA methods to infer landscape parameters [21, 22]. While these approaches were effective at determining contacting residues in the three-dimensional structure, they ignore the sequential nature of laboratory evolution data and instead treat sequences from multiple generations as independent samples. Observing how the evolutionary process unfolds over time provides valuable information about the structure of the fitness landscape.

In this work, we develop a statistical learning framework to infer the protein fitness landscape from laboratory evolution data. We use population genetics principles to develop a model of the underlying evolutionary process and build a likelihood function to estimate the landscape parameters from multiple rounds of evolution. We performed 15 rounds of laboratory evolution on the enzyme dihydrofolate reductase (DHFR) to generate a large and diverse data set consisting of sequences sampled from multiple sequential generations. We applied our learning method to infer the DHFR landscape and found the model parameters capture important aspects of DHFR function and reveal landscape epistasis arising from interactions between residues. We used the learned model to understand the global structure of the fitness landscape by running thousands of evolution simulations and found all trajectories converged to the same sequence, suggesting a single global optimum despite many examples of local epistasis. Finally, we applied our model to start from where our experimental DHFR evolution left off and continue the evolutionary process *in silico*. This procedure was used to extrapolate the evolutionary trajectory and design new functional DHFRs that were beyond the training data.

## Results

### Laboratory evolution to explore the fitness landscape of dihydrofolate reductase

Directed laboratory evolution applies iterative rounds of mutation and selection to explore the protein fitness landscape. We performed a directed evolution experiment on murine dihydrofolate reductase (mDHFR) to search the fitness landscape for diverse sequences encoding DHFR activity (Figure 1). DHFR reduces dihydrofolate to tetrahydrofolate and plays an essential role in purine biosynthesis and cell growth. We mutagenized mDHFR using error-prone PCR, transformed the library into *E. coli*, and performed a selection to identify DHFR variants capable of supporting *E. coli* growth. We then remutagenized all surviving variants, repeated the process, and performed a total of 15 rounds of directed evolution. Each round of evolution introduced an average of four DNA mutations into the DHFR genes and maintained a population size of 10^5^-10^6^ transformants. Importantly, our low stringency selection and large population size create a neutral evolutionary process that generates diverse sequences that maintain wild-type-like DHFR activity.

**Figure 1.**
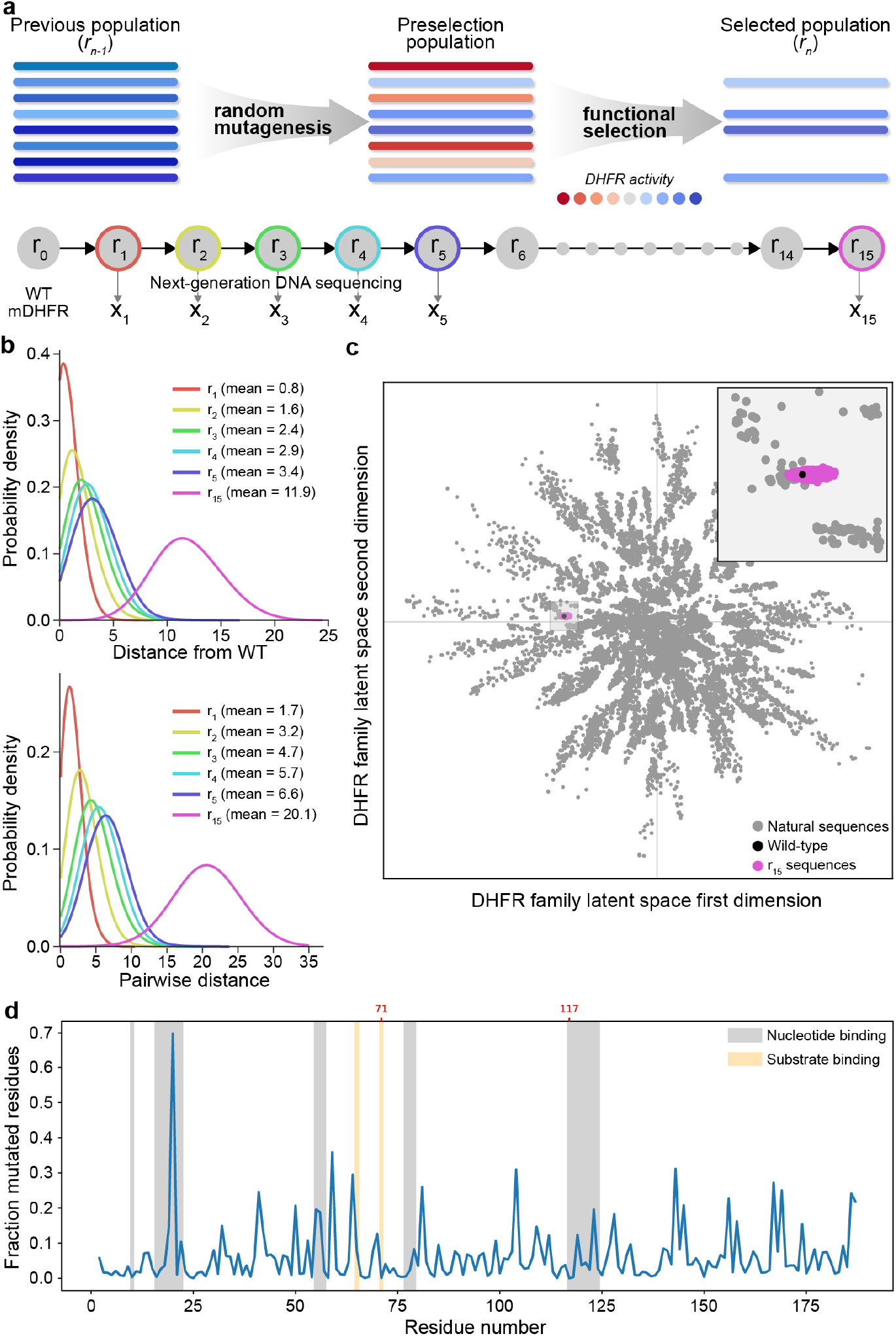
DHFR laboratory evolution (a) Directed laboratory evolution combines iterative rounds of random mutagenesis and functional selection to evolve populations of molecules. We performed 15 rounds of directed evolution on DHFR and sequenced the population at generations 1-5 and 15 to obtain snapshots along the evolution trajectory. (b) Sequence statistics of the evolving populations displayed increasing distance from wild-type DHFR and increasing distance between sequences within a population, suggesting a diffusion-like spread into protein sequence space. (c) Visualization of the round 15 population in the context of natural DHFR sequences. The laboratory evolution experiment explored a small fraction of DHFR sequence space. Sequences were visualized using a two dimensional latent space of a variational auto encoder (VAE) trained on natural DHFR sequences. (d) The mutational statistics of sequences from the round 15 population displayed lower mutation rates at active site residues. The mutation N20D became fixed in the round 15 population.

We performed Illumina DNA sequencing on directed evolution rounds 1-5 and 15 to obtain a sample of the evolutionary trajectory. From this data, we see the evolutionary process generates a distribution of sequences with varying Hamming distance from wild-type mDHFR and the evolving population increasingly drifts away from the starting sequence (Figure 1b). The round 15 population has an average of 11.9 amino acid substitutions, corresponding to 0.79 amino acid substitutions accumulated per round. We also observed the populations’ pairwise Hamming distance increased linearly over the course of evolution. The fact that the average distance from wild-type and the average pairwise distance both increased linearly with each round indicates that most evolutionary trajectories are exploring independent directions on the landscape. We also visualized how the DHFR sequences generated in our directed evolution experiment fit into the larger protein family generated by natural evolution (Figure 1c). Our directed evolution experiment started to capture similar variation to some natural sequences but only explored a small fraction of the sequence space spanned by nature.

We further analyzed the round 15 sequences to understand how mutations were distributed across the primary sequence. The mutations generally are distributed across the protein sequence but there is a lower observed mutation rate in the protein core and active site residues. We also observed the N20D mutation overtook the wild-type asparagine residue in the round 15 population, possibly indicating positive selection for this mutation. Residue 20 is in the active site region of mDHFR that binds nucleotide phosphates.

### A statistical framework to learn from sequential rounds of experimental evolution

Directed evolution provides a sampling of populations evolving on a protein fitness landscape. We develop a statistical framework to infer the underlying landscape structure from these experimental evolutionary trajectories. We posit that data from sequential rounds of evolution provides inherently more information than data from a single round or data from multiple rounds considered independently. Observing how the evolutionary process unfolds over multiple rounds allows us to make stronger inferences and extrapolate behaviour. For simplified illustrative example, if we observe how the frequencies of amino acids change over each round, we can extrapolate these trajectories to estimate where the evolutionary process will converge. We build a generative model of the laboratory evolution process, parameterize the fitness landscape using a generalized Potts model, and infer the landscape parameters from sequencing data from multiple rounds of evolution (Figure S2).

We model the dynamics of laboratory evolution as a Markov chain process where sequences transition to other sequences according to their mutational accessibility and relative fitness. We consider the set of all possible codon sequences Ω of length *L*, where each sequence is denoted by *x* ≔ (*x*_1_, …, *x_L_*). Each *x_i_* corresponds to the codon that encodes the *i*th residue position and each codon is from the set 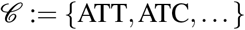 of 61 codons that exclude the stop codons. We let Π(*x*) represent the fitness of each sequence *x* ∈ Ω, i.e., the number of copies that a sequence makes of itself per *unit* time, and *π*(*x*) the corresponding prevalence based on relative fitness, defined as *π*(*x*) = Π(*x*)/ ∑_*u*∈Ω_ Π(*u*).

We make a number of assumptions on the transition mechanism between rounds to model the laboratory evolution with a Markov chain process. First, we assume the mutation process *g* happens independently at each DNA position and the mutation probability at each position is known from the experiment. Second, we assume that the transition mechanism is time-homogeneous, that is, the both fitness values and the mutation probabilities do not change between rounds. Third, we assume that number of transformants is sufficiently large so that the distribution of sequences at round *r* only depends on the wild-type sequence and the relative fitness level *π*. Finally, we assume simplified Markov chain dynamics, which assumes localized competitions between direct descendants, well approximates the true dynamics (see Supplementary, Sec. Markov chain approximation to infinite population dynamics for details). Under these assumptions, we model the dynamics of laboratory evolution as a Markov chain process where the chain starts at the wild-type sequence and the transition probability from sequence *x* in round *r* to sequence *y* in round *r* + 1 is given by

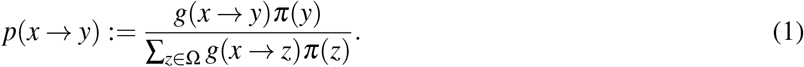

where *g*(*x* → *y*) represents the probability sequence *x* mutating to sequence *y* in the absence of selection. Each sequence *x*^(*n,r*)^ from the round *r* is then a random sample from the marginal probability *p*^(*r*)^ after the *r*th step transition.

We parameterize the fitness landscape *π* with a (generalized) Potts model that describes how all amino acid residues and their pairwise interactions contribute to fitness. Potts models have been used extensively to recover the interaction graph between residues of a protein and strong interactions have been shown to correspond to long-range contacts in the 3D structure of a protein [15, 17]. We parameterize the fitness level *π*(*x*) = *π_θ_* (*x*) of sequence *x* with a Potts model with canonical parameter set on the amino acids 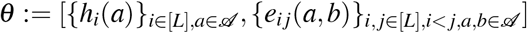 where

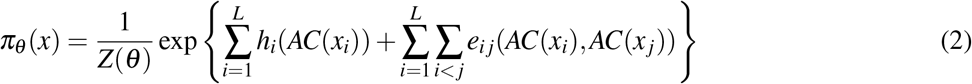

AC(·) is the mapping from the set of codons 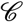 to the set to amino acids 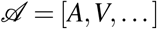, and *Z*(*θ*) is the normalization constant so that the probabilities sum to 1. This model consists of *q_a_L* main effect parameters (*h_i_*), where 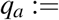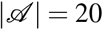, and 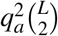 (couplings) interaction effect parameters (*e_i j_*).

Directed evolution data consists of sequences sampled from multiple rounds of evolution. These observed sequences *x*^(*n,r*)^, *n* = 1, 2, …, *n_r_* at each round *r* are random realizations from the probability distribution *p*^(*r*)^(·; *θ*). We can estimate the Potts model parameters *θ* by maximizing the following log-likelihood

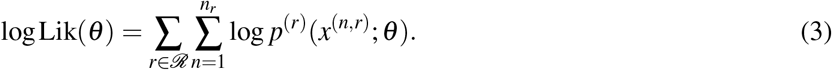

where ℛ denotes the set of experimental rounds at which sequencing data exists and *p*(*x* → *y*; *θ*) in (1) is given by the Markov chain dynamics under the Potts model.

It is challenging to identify the model parameters *θ* that maximize this log-likelihood function (3). First, even though *π_θ_* forms a Markov Random Field (MRF), the distribution of *p*^(*r*)^ no longer factorizes with respect to the graph associated with the fitness distribution *π_θ_*. In particular, the conditional independence relationships which hold for *π_θ_* do not hold for *p*^(*r*)^, *r* = 1, 2, …, which prohibits applying techniques for parametric inference in Markov Random Field settings. In addition, the large dimension of the state space precludes any exact tracing of this Markov chain process. For instance, the dimension of the transition matrix is |Ω| × |Ω|, and even computing a single element in the transition matrix for a given *θ* is computationally infeasible due to the intractable sum in the denominator in (1) (as |Ω| = 61^186^).

We used an approximate moment matching method to overcome these computational challenges. In particular, we first derived approximate relationships between the first and second order marginals 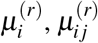 at each round *r* and the marginals *μ* under *π_θ_*. Then we sought for 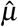 which matches empirical and expected 1st and 2nd order marginals and is also locally compatible (Supplementary, Sec. Inferring mean parameters). Lastly, we used mean-field DCA [15] using estimated marginals as inputs to get estimates of parameters of the Potts model (2). The details of the statistical method are available in (Supplementary, Sec. Overview of statistical method).

### Learned landscape parameters capture protein structure and function

We applied the statistical learning framework developed above to infer DHFR’s fitness landscape from our experimental laboratory evolution data. We estimated the Potts model’s canonical parameters from evolution rounds 1-5 and 15, and used these parameters to get information about the protein’s structure and function.

The learned Potts model reveals how individual amino acid substitutions affect wild-type DHFR’s activity (Figure 2a). This learned mutation map clearly highlights the importance of the enzyme’s key catalytic residues and displays expected mutational patterns residues with similar physiochemical properties. The G18A amino acid substitution has the largest beneficial effect and is located in a loop that interacts with NADPH. We mapped the average mutational effect magnitude onto the three-dimensional DHFR structure to understand how the learned parameters relate to structure (Figure 2b). We observed that mutations in the protein core tend to have larger effects on activity, presumably because mutations at these sites disrupt the enzyme’s three-dimensional structure.

**Figure 2.**
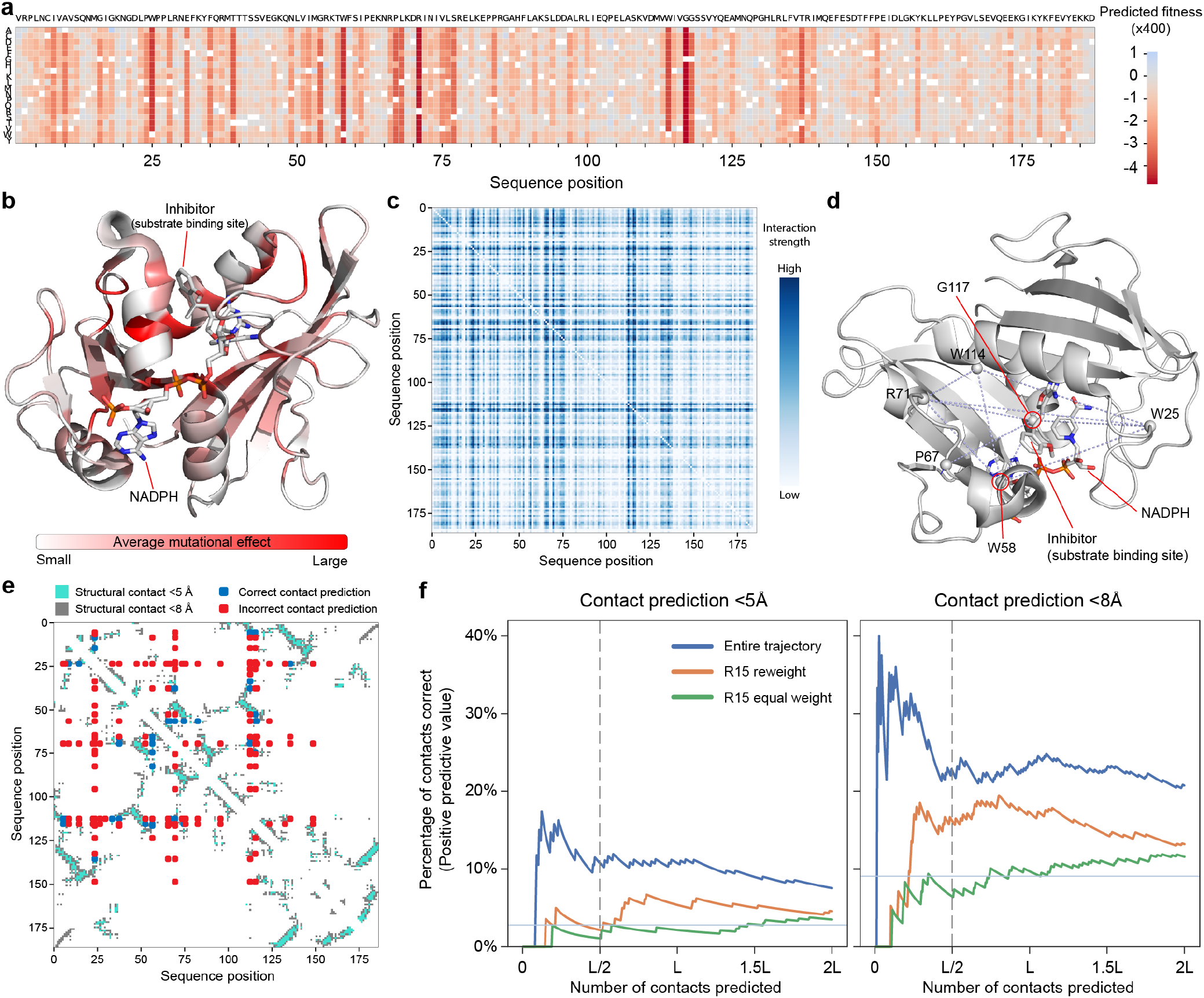
Model parameters relate to DHFR structure and function. (a) A heatmap of the model’s predicted mutational effects across the 186 DHFR sequence positions. The wild-type amino acid is colored in white. (b) The average mutational effect at each site mapped onto the DHFR structure (PDB ID: 3K47). The largest magnitude mutations tend to occur in the protein core and the substrate binding site. (c) The interaction strength between all pairs of sites in DHFR. The interaction strength is calculated as the Frobenius norm over the Potts model interaction coefficient all amino acid combinations at each pair of sites. (d) The top ten long range interactions plotted on the DHFR structure (PDB ID: 3K47). Many of these interactions occur through interactions through the substrate. (e) The top L/2 (93) interactions between residues plotted on a contact map showing residues with heavy atoms closer than 5 Å and 8 Å. (f) A comparison of contact prediction for pseudo-likelihood DCA models trained on R15 data, R15 data reweighted to account for evolutionary bias, or using our methods on the entire evolutionary trajectory. The horizontal line represents random chance and the vertical line is drawn at the commonly used *L*/2 threshold.

The Potts model can also be used to understand landscape epistasis and interactions between residues. We identified only two examples of reciprocal sign epistasis in the landscape out of approximately 6 million possible amino acid and position pairs. One such example occurs between A47K and R99E, where the individual mutations are beneficial individually, but when combined result in a decrease in fitness. We observed 871,296 examples of sign epistasis in the landscape where the double mutant and its two single mutants do not all have the same sign increase or decrease in fitness. We computed a residue-residue interaction score by calculating the Frobenius norm *F_i j_* between all interaction parameters between a pair of residues (Figure 2c). The inferred residue interactions capture many contacts from the enzyme’s three-dimensional structure and also functional interactions in the enzyme active site. The top interaction is between residues R71 and G117, which are not directly interacting in the 3D structure, but form opposite ends of the nucleotide binding pocket (Figure 2d). Seven of the top ten residue-residue interactions involve one of these key sites. The top 20 interactions are provided in Table S1.

The Potts model interaction scores can be used to identify residue pairs that are contacting in the three-dimensional protein structure. The residue pairs with the top *L*/2 (93) Frobenius scores correspond to 28 structural contacts at a distance of less than 5 Å and 77 contacts at a distance of less than 8 Å (Figure 2e). We compared our model’s contact prediction performance to established methods for inferring contacts from evolutionary data, including the standard DCA modeling procedure [19] and also a modified DCA model that weights sequences based on their distance from wild-type to account for the evolutionary process [22] (Figure 2f, Table S2). When trained on the round 15 data, neither of these methods were able to outperform the random chance expectation (2.79%) when predicting the top *L*/2 (93) long range contacts at a distance of less than 5 Å. In contrast, our model, which considered the entire evolutionary trajectory, was able to correctly identify 10 out of the top *L*/2 contacts (10.7%), which is well above random chance. Our model also outperforms the other two methods by recovering more structural contacts with heavy atoms 5 − 8 Å and less than 8 Å (Figure 2f, Table S2).

### In silico evolutionary simulations to map the global structure of the fitness landscape and extrapolate evolutionary trajectories

Our statistical method infers the underlying fitness landscape from experimental laboratory evolution trajectories. This model can be used to run *in silico* simulations to understand the landscape, evolutionary processes, and engineer new proteins.

We used our model to understand the global structure of the fitness landscape and the convergence of adaptive evolutionary walks. We sampled 1600 random amino acid sequences to obtain broad sampling of the landscape. For each of these sequences, we performed an adaptive walk by evaluating all single mutants, selecting the most fit variant, and repeating this process until a local fitness peak was reached. We found every single adaptive evolutionary trajectory converged to the same fitness peak and the sequence at this fitness peak is 79 amino acid substitutions from wild-type DHFR. The fact that adaptive walks starting from diverse regions of the landscape converge to the same peak implies a Mt. Fuji-type fitness landscape with few local optima.

We also used the learned model to continue the experimental DHFR evolution process and extrapolate the evolutionary trajectory to future generations (Figure 3b). We started the simulation with the most common amino acid sequence observed in the final round of the laboratory evolution experiment. This starting sequence is 10 amino acid substitutions from wild-type DHFR. We performed an adaptive walk by evaluating all single and double mutants, selecting the most fit variant, and repeating this process until there were no further uphill steps. The simulated evolutionary trajectory continued to move away from wild-type DHFR and the round 15 starting sequence and converged to the same fitness peak as found in the global landscape search described above.

**Figure 3.**
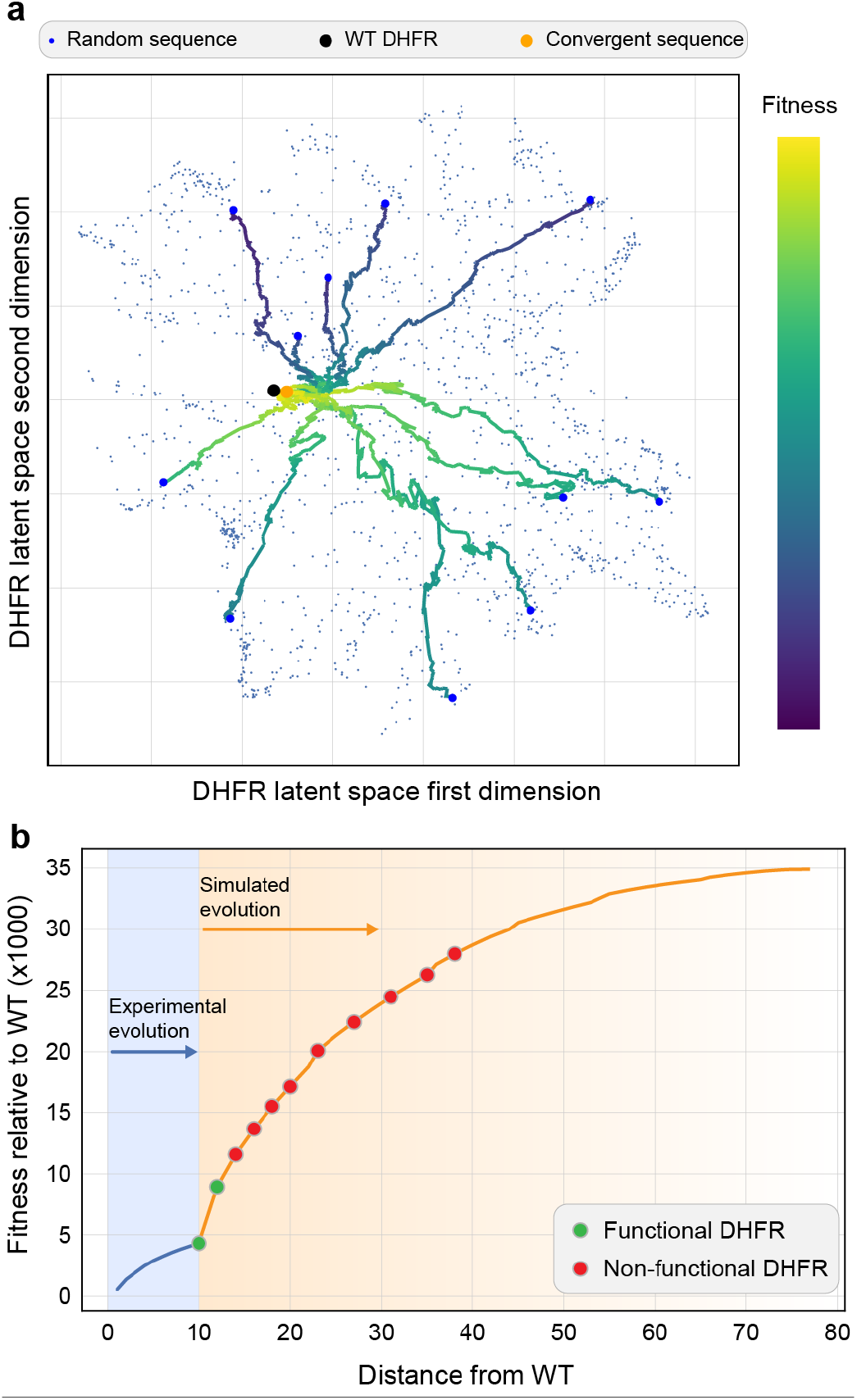
Evolutionary simulations to understand the landscape structure. (a) We sampled random sequences from a 2D VAE latent space and performed an adaptive walk until a fitness peak was reached. All adaptive walks converged to the same fitness peak. 11 representative evolutionary trajectories are shown. (b) We continued our laboratory DHFR evolution by continuing the evolutionary process *in silico*. We experimentally tested several DHFR sequences along the evolutionary trajectory and found many were inactive enzymes.

We wanted to test whether the directed evolution trajectories could be extrapolated *in silico* as an approach to engineer new proteins. We experimentally characterized ten DHFR sequences along the trajectory from the round 15 sequence to the adaptive fitness peak (Figure 3b). We found the round 15 sequence was an active DHFR enzyme, a double mutant of the round 15 sequence was also active, but all sequences beyond that were inactive and unable to complement *E. coli* growth in the presence of trimethoprim (Figure 3b, Figure S3).

## Discussion

Directed laboratory evolution is a powerful approach to explore the protein fitness landscape. Directed evolution generates data consisting of sequences sampled over multiple generations and provides valuable clues about the underlying fitness landscape structure. Directed evolution data does not naturally fit into established supervised or unsupervised learning methods because they do not consider the evolutionary data generation process. In this work, we developed a statistical learning framework to infer the protein fitness landscape from laboratory evolution data. We built a generative model of the directed evolution process, parameterized the fitness landscape using a generalized Potts model, and inferred the landscape parameters from sequencing data from multiple rounds of evolution. We applied this learning approach to a large and diverse dihydrofolate reductase (DHFR) directed evolution data set. The inferred landscape model revealed numerous examples of epistasis arising from interactions between residues, but an overall global fitness peak that is evolutionarily accessible from most starting sequences. Finally, we explored the potential of the landscape model to extrapolate evolutionary trajectories for protein engineering.

Our laboratory evolution model made two important approximations for the sake of computational tractability. Understanding when these approximations are valid or invalid can help identify the scope and limitations of our method. The first approximation makes the assumption the evolving population has an infinite population size, while in practice all laboratory evolution has a finite population size dictated by experimental constraints. Our actual DHFR experimental population sizes are on the order of 10^5^ – 10^6^ and this number is largely determined by the library transformation efficiency into E. coli. With larger population sizes, this approximation gets more accurate.

The second approximation makes the assumption that the population evolves according to simplified Markov chain dynamics (1) where competition only occurs between sequences who share a direct ancestor. In our actual experiment all sequences are directly competing in the growth selection regardless of their lineage (Figure S1). The Markov chain dynamics approximation is valid when competing sequences have similar fitness at each stage of the experiment, which is likely if the fitness landscape near the initial wild-type sequence is flat (i.e., most selected mutations are neutral) and the evolution experiment is carried out at lower mutation rates. At low mutation rates the laboratory evolution experiment only explores the wild-type sequence’s local neighborhood and all selected mutant sequences will have similar fitness due to the neutral landscape. In addition, the Markov chain dynamics are a better approximation during the earlier rounds of evolution, while all evolving sequences are still near the initial wild-type sequences.

Once we specified the evolution dynamics, we inferred fitness landscape parameters by approximate moment matching. The approximate recursive formulas for the first and second order moments between round *r* and *r* + 1 were derived based on a weak dependence assumption between variables of interest with the remaining variables, which allow us to approximate marginals at *r* + 1 round only based on local information in the protein residue graph. This approximation is only exact if the residue graph factorizes, but we expect this provides a reasonable approximation for most of the nodes and/or pair of nodes where the strength of interaction between them and the other variables is limited. We have three main regularization hyperparameters in our inference procedure. The first two influence the magnitude of the main effects and the pairwise effects parameters. These can be set in the same way as they are for the DCA model [17]. The third hyperparameter is the regularization parameter that infers the canonical parameters from the mean parameters. We observed the best performance when it was set to be several times larger than the negative of the smallest eigenvalue in the estimated covariance matrix. Of these three parameters, the results were most sensitive to the last parameter.

Our inferred DHFR landscape parameters were able to identify contacting residues in the three-dimensional protein structure and also numerous functionally coupled residues that indirectly interact through the enzyme active site. Our trajectory learning approach showed significant improvements over established contact prediction methods at the commonly used *L*/2 threshold for residues with heavy atoms closer than 5 Å. This improvement is due to modeling the evolutionary trajectory as well as including sequencing data from earlier rounds of evolution. Several of top interactions inferred are in residues that are not close in the 3D structure but are in functional regions that interact with either the substrate or the cofactor. These interactions are marked as false positives for contact prediction, however they are likely epistatic interactions arising through ligand interactions or cofactor repositioning [23].

Previous work has demonstrated the ability to identify residue contacts by applying DCA to the final generation of laboratory evolution experiments [21, 22]. Stiffler et al. applied a reweighing procedure to DCA to account for evolutionary bias and were able to get contact recovery rates exceeding 50% for the top *L*/2 contacts. Previous laboratory experiments using TEM1 *β*-lactamase [24] were not able to detect local epistatic interactions, however a larger and more recent experiment [21] was able to identify epistatic interactions and detect some contacts using DCA. It’s notable that these DCA methods were unable to reliably detect DHFR contacts from our round 15 evolution data (Figure S4). We applied standard DCA and the reweighting DCA methods of [22] to our round 15 DHFR data and when predicting the top *L*/2 contacts closer than 5 Å in the 3D structure, the percentage of correctly labeled contacts was lower than random chance expectation. There are a number of factors that could contribute to the differences in contact recovery across these data sets, including population sequence diversity, the number of variants sequenced, or the evolution mutation rate. Stiffler et al. provide an analysis of down-sampled data sets for aminoglycoside acetyl-transferase (AAC6) [22, Figure 3A], so we can directly compare to our mDHFR data. At 10^5^ sequences, they have an average pairwise distance of 10.9% and identify nearly 40% of the contacts. In contrast, with our mDHFR data with 10^5^ sequences, we have an average pairwise distance of 11.1%, but only recover 2.15% of the contacts. Based on these findings, it seems the sequence diversity and data set size are not contributing to differences in contact recovery. The two experiments do differ significantly in their mutation rate, where AAC6 0.8% per round, while DHFR was evolved at 0.4% per round. We hypothesize this varied mutation rate is resulting in different population structures, despite having similar sizes and levels of diversity. Even for a fixed neutral landscape, finite populations following the quasispecies dynamics with different mutation rates will evolve different levels of robustness [25] and this robustness is linked to epistatic interactions [26]. Simulations of these types of laboratory evolution experiments using the simple markov chain dynamics is provided by [27] and these simulations do not consider the mutation rate as an important parameter. However, since the laboratory evolution experiment likely follows the more complicated quasispecies dynamics, it is possible that the mutation rate plays a more important role.

We used the inferred model parameters to explore epistasis and the global structure of the fitness landscape. We found many examples of additive mutations, approximately 10% of mutations that interact through sign epistasis, and only two examples of reciprocal sign epistasis. This frequency of epistatic interactions is consistent with other studies [28, 29]. Despite the landscape epistasis, we found the estimated landscape structure had a overall global fitness peak that was accessible by adaptive walks from most starting sequences. Similar fitness landscape features have been observed in other proteins [29, 30].

We used the inferred Potts model to run an *in silico* directed evolution experiment to design new, previously unobserved DHFR variants. We found the model could design functional DHFRs that were close to the training data regime, but further evolutionary extrapolation resulted in nonfunctional enzymes. This result is consistent with other ML-based protein engineering studies that show decreased model accuracy while extrapolating away from the training data [10, 31]. The model inaccuracy could be the result of insufficient or low-quality data, approximations we made in the evolutionary model, or computational challenges with parameter estimation. A more accurate evolutionary model would consider finite population sizes and competition between all sequences in a generation, which is referred to as the finite quasispecies model [25, 32]. Another possibility to improve model accuracy would be to use a simpler first order model without interactions so that we need to estimate fewer parameters. Another approach to improve the reliability for protein engineering would be to run multiple independent evolution simulations and test a panel of diverse designs.

Our statistical landscape inference approach naturally complements recent advances in continuous directed evolution [33–35]. These experimental methods combine population level mutagenesis and selection in continuously-fed bioreactors to evolve populations without discrete mutation/selection steps. The populations can be sampled and analyzed by next-generation DNA sequencing to observe how the population changes over time and traverses the fitness landscape. Our learning method could infer the landscape from this sequential evolution data to understand protein structure, function, and evolution.

The relationships between protein sequence, structure, and function involve thousands of exquisite molecular interactions that are dynamically coupled over space and time. Machine learning is revolutionizing our understanding of these relationships by dissecting the complex inner workings of proteins with a scale and resolution beyond human comprehension. Future advances in data-driven protein science will improve our ability to understand natural evolutionary processes, predict genetic disease, and design new proteins for broad applications in biotechnology.

## Materials and methods

### mDHFR laboratory evolution

Our selection strain consists of the murine dihydrofolate reductase (mDHFR) gene cloned into the pET22b plasmid and transformed into *E. coli* BL21(DE3). We performed error-prone PCR using the reaction’s MnCl_2_ concentration to tune the mutation rate of Taq DNA polymerase [36]. We determined that a final concentration of 200 *μ*M MnCl_2_ yielded 3.25±0.74 amino acid substitutions per gene. We performed 15 error-prone PCR cycles, treated the reaction with DpnI overnight to remove template, purified the PCR product with a DNA spin column (Zymo Research), cloned the insert back into pET-22b using circular polymerase extension cloning (CPEC) [37], purified the CPEC reaction using a DNA spin column (Zymo Research), and transformed the CPEC reaction into electrocompetent BL21(DE3) cells (Lucigen). Several dilutions of the transformation were plated to determine the total library size, which was in the range of 10^5^ – 10^6^ colony forming units (CFUs). The remainder of the transformation was used as input to a competitive growth selection in 100 mL of LB containing 100 *μ*g/mL carbenicillin, 500 *μ*M IPTG, and 5 *μ*g/mL trimethoprim. We allowed these selection cultures to grow shaking for 16 hours at 37 °C. Approximately 20 ODU of overnight culture was used to harvest plasmid DNA via miniprep. This selected plasmid DNA population was then used as a template for the next round of error-prone PCR. A portion of the post-selection culture was also archived as 15% glycerol stocks and stored at −80 °C.

We determined the fraction of functional variants from each round of evolution by picking colonies from each transformation plate into individual wells of a 96-well plate containing LB broth, 100 *μ*g/mL carbenicillin, 500 *μ*M IPTG, and 5 *μ*g/mL trimethoprim. We incubated these plate cultures shaking for 16 hours at 37 °C, measured the OD600 of each well, and categorized DHFR variants as functional if their OD600 was greater than 0.5, otherwise they were considered nonfunctional.

### DNA sequencing of evolved populations

We performed next-generation DNA sequencing on several rounds of laboratory evolution. We analyzed rounds 1-5 using Illumina sequencing and round 15 using Pacific Biosciences sequencing. For the Illumina libraries, we used NdeI and SacI restriction enzymes to remove the DHFR insert from the plasmid, ligated Illumina adaptor sequences to this insert, and submitted the samples to the UW-Madison Biotechnology Center DNA Sequencing Core to run on an Illumina MiSeq instrument using the 2×300 v3 kit. Each sample had 2-5 million reads. For the PacBio sequencing, we removed the DHFR insert with NdeI/SacI and submitted the samples to the UW-Madison Biotechnology Center DNA Sequencing Core for analysis on their Pacific Biosciences Sequel instrument. The PacBio run returned over 10^5^ reads.

### Sequence pre processing

For rounds 1-5, the Illumina sequencing data are processed with steps similar to [22]. The forward and reverse reads are first stitched together using the FLASH program [38]. At the first filtering step, only sequences with a minimum length of 500 and a minimum quality score of 15 for every base were retained. At the second step, sequences with a compound quality score of atleast 10 were retained, implying a 90% probability of having no read errors. For round 15, quality filtering was done to keep sequences with a minimum length of 564 and a minimum quality score of 0.99. The remaining sequences were then aligned to the reference sequence using bowtie2 [39].

### Statistical framework parameters

The statistical inference method was implemented in Python using PyTorch [40]. The optimization method used the Adam optimizer with a learning rate of 0.03 with other learning parameters set to default and trained for 300 steps. The regularization hyperparameters in (Supplementary, Eq. 14) were set to *λ*_main_ = 10^−3^, *λ*_int_ = 10^−4^ and *λ*_reg_ = 50. The penalty term to ensure the main affect parameters marginalized to the pairwise parameters was set to *ρ* = 10^5^.

The exact mutation distribution in this experiment is not known as all rounds were sequenced after growth selection, however, the statistical method seems fairly robust to choice of the mutation distribution as the results look similar with mutation bias distributions from other experiments done with a similar protocol (results not shown). We picked a mutation bias distribution to model mutagenesis from the *Taq* DNA polymerase column of [41, Table 2], [42] and then scaled to match an average of 4 DNA mutations per round. The final mutation distribution used in the statistical method is given in Table S3.

### Experimentally testing evolutionary designed DHFRs

We designed ten DHFR variants using the inferred model to simulate an evolutionary process. The genes encoding these variants were synthesized by Twist Bioscience and cloned into the pET21(+) plasmid. We transformed the plasmids into *E. coli* BL21(DE3) and performed growth measurements to assess the DHFR variant’s activity. We performed the growth assays by first inoculating a 5 ml LB starter culture containing 100 *μ*g/mL carbenicillin and growing overnight shaking at 37 °C. We then diluted this starter culture 100x into an LB culture containing 100 *μ*g/mL carbenicillin, 500 *μ*M IPTG, and 5 *μ*g/mL trimethoprim and monitored growth by measuring the OD600 in 30 min. intervals over a 16.5 hour incubation period at 37 °C. These measurements were carried out in triplicate. Inactive DHFR variants displayed no growth under these conditions, while active variants displayed standard growth curves.

