## Supplement for "Inferring protein fitness landscapes from laboratory evolution experiments"

### Supplementary Figures

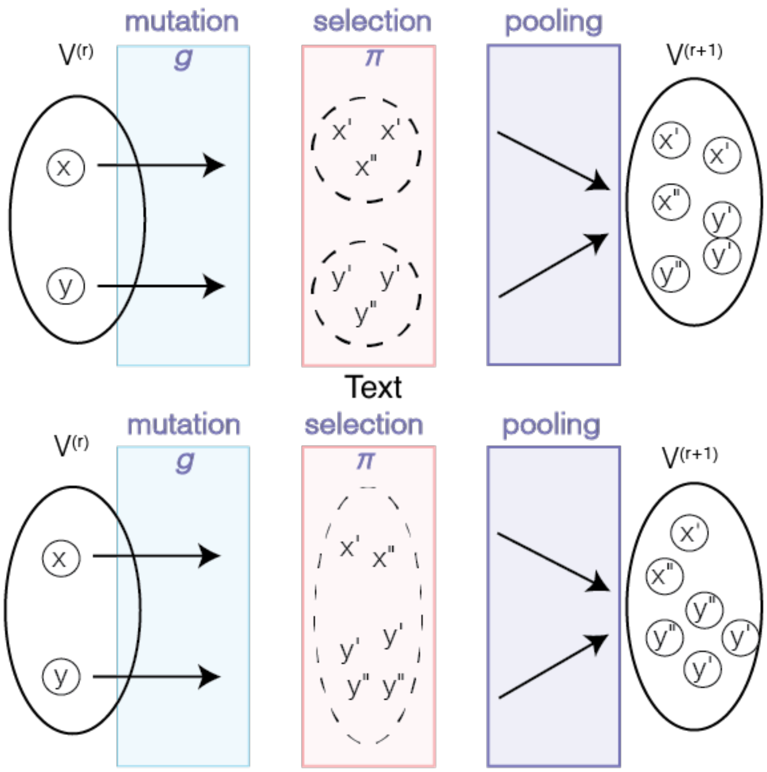


**Figure S1.** Diagram of an Idealized Experiment (first row) vs Real Experiment (second row). Suppose an overall capacity in the selection step is 6, and the relative fitness level of sequences $x', x'', y', y''$ is $x'' < x' \ll y'' < y'$. We see more $x', x''$ are present in $V^{(r+1)}$ in the first experiment due to a localized competition in $(x', x'')$ and $(y', y'')$.

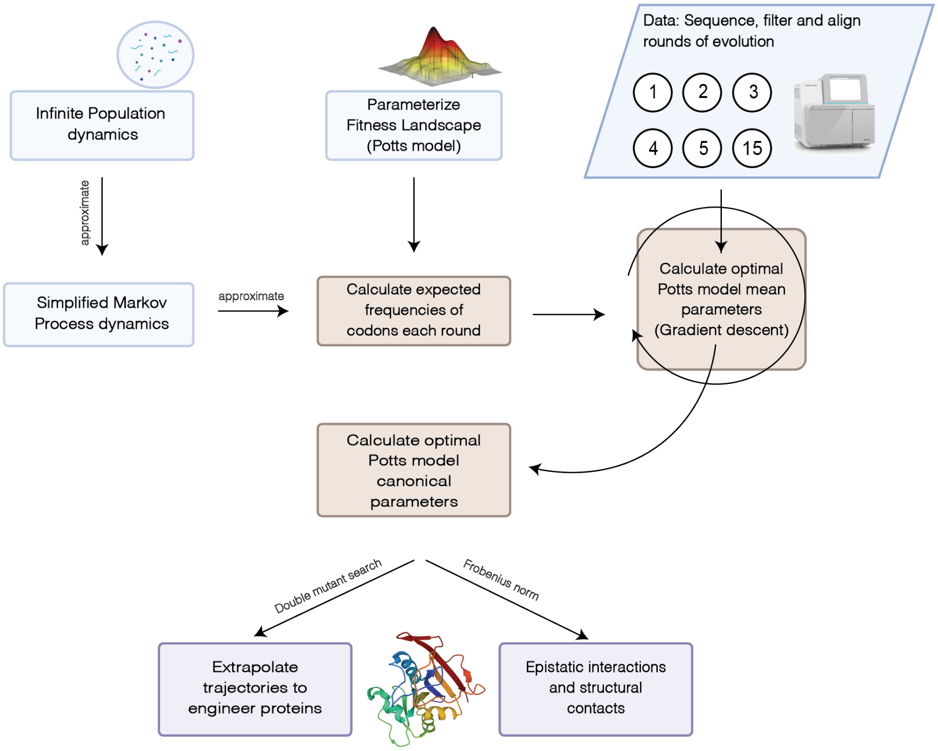


**Figure S2.** An overview of the algorithm to infer fitness landscape parameters.

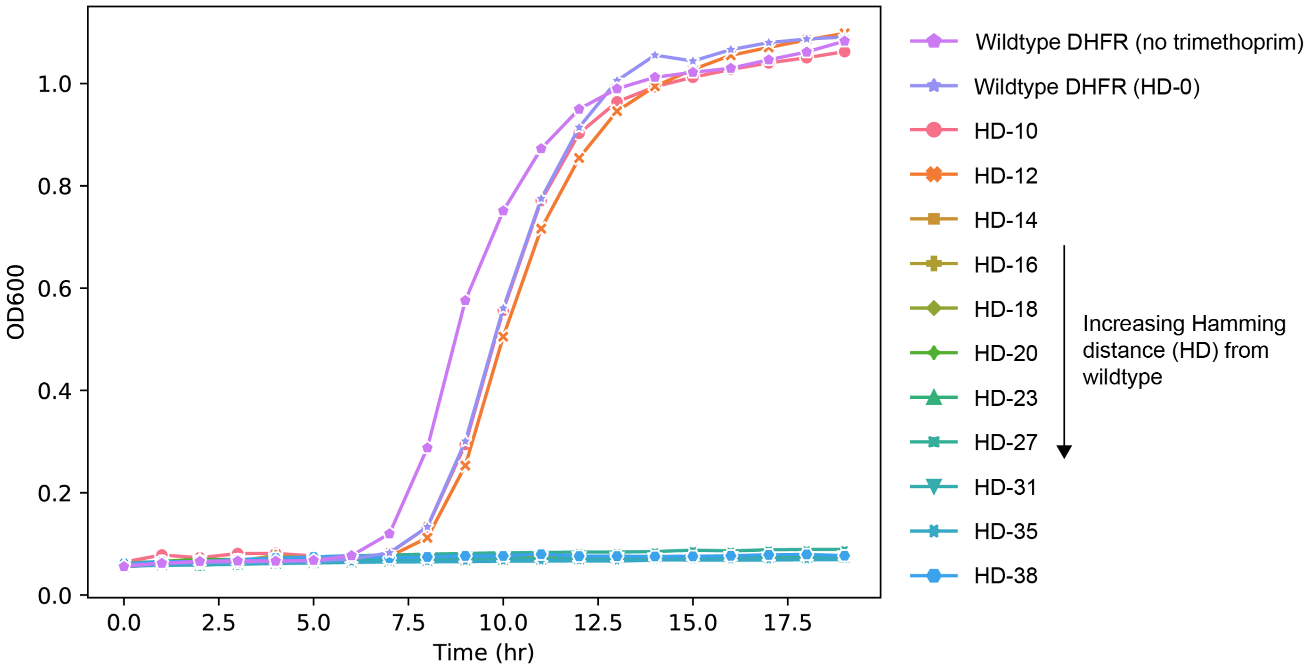


**Figure S3.** Growth curves of sequences designed using the inferred model, the most common sequence in round 15 and the wild-type sequence mDHFR.

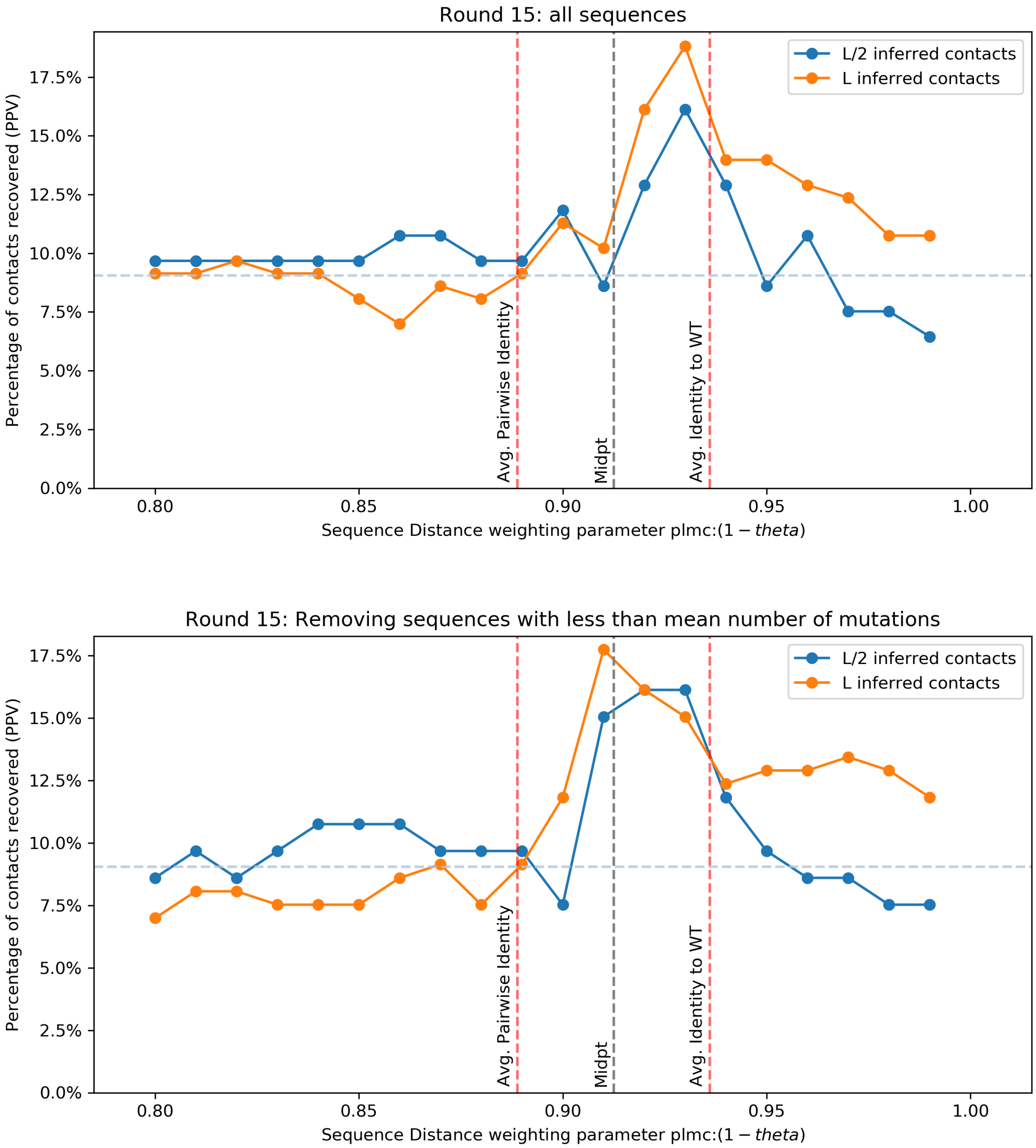


**Figure S4.** DCA methods trained on round 15 DHFR evolution data with different sequence weighting parameters (1 − theta). We ran *plmc* [1] on all sequences (top panel). Also, we follow the reweighting method of [2] (bottom panel). We first remove sequences at a distance less than the average distance from wild-type. Then we find that setting the sequence weighting parameter to the midpoint of the pairwise identity and the average identity to wild-type results is optimal in that it results in the highest positive predicted value when looking at the top $L$ contact predictions and close to optimal when looking at the top $L/2$ predictions.

### Supplementary Tables

**Table S1.** The top 20 long range interaction scores between pairs of residues. The columns AA1 and AA2 refer to the wild-type amino acid at the corresponding residue. We label the distance between the residues in the 3D structure. We also label whether the residue is involved in the function of the protein by labeling nucleotide binding regions or an isolated nucleotide binding site of the protein. Many of the top 20 interactions involve functional residues.

| | Res1 | Res2 | AA1 | AA2 | Residue distance | Frobenius Score | Contact | Res1 Function | Res2 Function |
|---|---|---|---|---|---|---|---|---|---|
| 1 | 71 | 117 | R | G | 46 | 0.000208 | no contact | Binding Site | Binding Region |
| 2 | 25 | 71 | W | R | 46 | 0.000175 | no contact | None | Binding Site |
| 3 | 71 | 114 | R | W | 43 | 0.000164 | 5-8Å | Binding Site | None |
| 4 | 25 | 117 | W | G | 92 | 0.000162 | no contact | None | Binding Region |
| 5 | 58 | 71 | W | R | 13 | 0.000157 | 5-8Å | None | Binding Site |
| 6 | 25 | 114 | W | W | 89 | 0.000138 | no contact | None | None |
| 7 | 25 | 58 | W | W | 33 | 0.000131 | no contact | None | None |
| 8 | 58 | 117 | W | G | 59 | 0.000127 | 5-8Å | None | Binding Region |
| 9 | 58 | 114 | W | W | 56 | 0.000123 | no contact | None | None |
| 10 | 67 | 117 | P | G | 50 | 0.000120 | no contact | None | Binding Region |
| 11 | 25 | 67 | W | P | 42 | 0.000118 | no contact | None | None |
| 12 | 67 | 114 | P | W | 47 | 0.000110 | no contact | None | None |
| 13 | 25 | 39 | W | T | 14 | 0.000109 | no contact | None | None |
| 14 | 25 | 118 | W | G | 93 | 0.000102 | no contact | None | Binding Region |
| 15 | 25 | 137 | W | T | 112 | 0.000102 | 5-8Å | None | None |
| 16 | 39 | 114 | T | W | 75 | 0.000102 | <5Å | None | None |
| 17 | 58 | 67 | W | P | 9 | 0.000102 | <5Å | None | None |
| 18 | 25 | 68 | W | L | 43 | 0.000099 | no contact | None | None |
| 19 | 25 | 54 | W | G | 29 | 0.000098 | no contact | None | None |
| 20 | 39 | 71 | T | R | 32 | 0.000095 | <5Å | None | Binding Site |

**Table S2.** Positive predictive value (PPV) of long range contacts recovered by different methods. These results are a table version of Figure 2f). $L = 186$

| Number of predictions | Method | <5Å | 5-8Å | <8Å |
|---|---|---|---|---|
| L/5 | Round 15 reweight | 2.70% | 2.70% | 5.41% |
| | Round 15 equal weight | 2.70% | 5.41% | 8.11% |
| | Entire trajectory (this paper) | 13.51% | 16.22% | 29.73% |
| L/2 | Round 15 reweight | 2.15% | 13.98% | 16.13% |
| | Round 15 equal weight | 1.08% | 5.38% | 6.45% |
| | Entire trajectory (this paper) | 10.75% | 11.83% | 22.58% |
| L | Round 15 reweight | 5.38% | 11.83% | 17.20% |
| | Round 15 equal weight | 2.15% | 7.53% | 9.68% |
| | Entire trajectory (this paper) | 10.75% | 12.90% | 23.66% |
| 2L | Round 15 reweight | 4.57% | 8.60% | 13.17% |
| | Round 15 equal weight | 3.49% | 8.06% | 11.56% |
| | Entire trajectory (this paper) | 7.53% | 13.17% | 20.70% |

**Table S3.** Error-prone PCR mutation bias. The transitions are given from columns to rows i.e. the probability of mutation from nucleotide $A$ to $C$ is given in column $A$ and row $C$.

| transition | A | C | G | T |
|---|---|---|---|---|
| A | 0.98860 | 0.00068 | 0.00205 | 0.00615 |
| C | 0.00110 | 0.99707 | 0.00021 | 0.00415 |
| G | 0.00415 | 0.00021 | 0.99707 | 0.00110 |
| T | 0.00615 | 0.00205 | 0.00068 | 0.98860 |

### Supplementary Methods and Mathematical Details

#### Markov chain approximation to infinite population dynamics

The dynamics of the prevalence (or concentration) of each sequence in each round are modeled as a mutation selection process and can be recursively written [3, 4] as

$$p^{(r+1)}(x) = \frac{\sum_{u\in\Omega} p^{(r)}(u)g(u\to x)\pi(x)}{\sum_{v,z\in\Omega} p^{(r)}(v)g(v\to z)\pi(z)} \tag{1}$$

The product in the numerator represents the number of sequences of $x$ produced by sequence $u$ as it is the product of the prevalence of sequence $u$ in round $r$, the mutation probability from sequence $u$ to $x$ and the fitness level of $x$. The denominator represents the total production of new sequences produced in round $r+1$ and it scales the prevalence vector $p^{(r+1)}$ so that it sums to 1. It involves the sum over all sequences that can possibly be mutated from *any* sequence that exists in the previous round. Since we start with only copies of the single wild-type sequence, the initial prevalence vector of sequences is 1 for the wild-type sequence $w$ and 0 for all other sequences. If we know the fitness values $\pi(x)$ and the mutation probabilities $g(u\to x)$, then Equation (1) completely determines the prevalence of every sequence $x\in\Omega$ in every round $r$. We call the dynamics represented by Equation (1) the **infinite population dynamics**.

The dynamics of Equation (1) are hard to work with even when the fitness levels $\pi$ are known. The equation (1) is not linear in the density $p^{(r)}(u)$, and the normalizing constant in the denominator involves a computationally intractable sum due to having to consider all sequences at the $r$th round as well as any possible sequence that can be mutated from sequences at the $r$th round. Facing such challenges, we propose to use approximate the infinite population dynamics in (1) via a linear dynamics which assumes competitions of fitness between sequences are localized among those which are mutated from the same direct ancestor. To illustrate, we consider an idealized experiment where for each sequence $x$ at round $r$, an isolated environment is provided where only sequences which are mutated from $x$ grow and compete in each environment during the $r+1$ round. At the end, all sequences are pooled together and the prevalence vector at the $r+1$th round $p^{(r+1)}$ is computed.

In this idealized experiment, the prevalence vector $p^{(r+1)}$ are given by

$$p^{(r+1)}(x) = \sum_{u\in\Omega} \frac{p^{(r)}(u)g(u\to x)\pi(x)}{\sum_{z\in\Omega} g(u\to z)\pi(z)}. \tag{2}$$

We can write this equation compactly in matrix form $p^{(r+1)} = S\,p^{(r)}$ where $S_{yx} := p(x\to y)$ where we recall that $p(x\to y)$ is defined as

$$p(x\to y) := \frac{g(x\to y)\pi(y)}{\sum_{z\in\Omega} g(x\to z)\pi(z)}. \tag{3}$$

We note $p^{(r+1)}$ is a valid probability vector because it is the product of a left stochastic matrix ($\sum_y S_{yx} = 1, \forall x$) and a stochastic column vector $p^{(r)}$. We call the dynamics represented by (3), **simplified Markov chain dynamics**, as we can think of it as arising from a Markov chain on the sequences of $\Omega$. The transition probability between sequence $x$ in round $r$ and sequence $y$ in round $r+1$ is given by $p(x\to y) := S_{yx}$.

The simplification from the infinite population dynamics (1) to the simplified Markov chain dynamics (2) is due to reducing many to many competitions to one to many competitions in the selection step (Figure S1), and therefore, the accuracy of approximation depends on abundance of sequences at each round as well as their relative fitness values. The approximation is expected to work well, for example, when there are relatively small number of distinct sequences as well as they share similar fitness properties, which is typically the case in neutral evolution experiment because introduced mutations tend to be neutral and a local neighborhood of the wild-type sequence is explored (Main, Fig. 1c). In the rest of the framework, we use the simplified Markov chain dynamics of (Main, Eq. 1) as an approximate model for the dynamics of the laboratory evolution experiment.

### Overview of statistical method

We first approximate first and second order marginals $\mu_i^{(r)}, \mu_{ij}^{(r)}$ with a function of first and second order marginals $\mu$ of $\pi_\theta$ (Sec. Approximate recursive relationships for marginals). Once the approximated moment vectors for all available rounds are obtained, the mean vector $\mu$ for $\pi_\theta$ is then estimated by matching the approximated moment vectors with the empirical counts.

We let $X^{(r)}$ be a random variable following the probability distribution $p^{(r)}$. $P(X^{(r)} = x) := p^{(r)}(x)$. For any codons $c, d \in \mathscr{C}$, first we define the first and second order marginals at round $r$ as $\mu_i^{(r)}(c) := \mathbb{E}[\delta(X_i^{(r)}, c); \theta], \mu_{ij}^{(r)}(c,d) := \mathbb{E}[\delta(X_i^{(r)}, c)\delta(X_j^{(r)}, d); \theta]$ where $\delta(\cdot,\cdot)$ is the Kronecker delta function such that $\delta(a,b) = 1$ if $a = b$ and 0 otherwise. We let the moment vector as a collection of these terms

$$\mu^{(r)} := [\{\mu_i^{(r)}(c)\}_{i\in[L],c\in\mathscr{C}}, \{\mu_{ij}^{(r)}(c,d)\}_{i,j\in[L],i<j,c,d\in\mathscr{C}}].$$

Similarly, we also define the first and second order marginals corresponding to the fitness landscape $\pi_\theta$ as $\mu_i(c) := \mathbb{E}_\pi[\delta(X_i, c); \theta]$, $\mu_{ij}(c,d) := \mathbb{E}_\pi[\delta(X_i, c)\delta(X_j, d); \theta]$, as well as the moment vector $\mu$ for $\pi_\theta$ as

$$\mu := [\{\mu_i(c)\}_{i\in[L],c\in\mathscr{C}}, \{\mu_{ij}(c,d)\}_{i,j\in[L],i<j,c,d\in\mathscr{C}}].$$

If we have sequencing data at round $r$, we can calculate the individual and pairwise codon frequencies observed for that round by summing over the data.

$$f_i^{(r)}(c) = \frac{1}{n_r}\sum_{n=1}^{n_r} \delta(x_i^{(n,r)}, c); \quad f_{ij}^{(r)}(c,d) = \frac{1}{n_r}\sum_{n=1}^{n_r} \delta(x_i^{(n,r)}, c)\delta(x_j^{(n,r)}, d)$$

We can put these terms together in a frequency vector $f^{(r)}$

$$f^{(r)} := [\{f_i^{(r)}(c)\}_{i\in[L],c\in\mathscr{C}}\{f_{ij}^{(r)}(c,d)\}_{i,j\in[L],i<j,c,d\in\mathscr{C}}\}],$$

and when it can be calculated from the data it gives us an estimate of the moment vector $\mu^{(r)}$ for round $r$.

The fitness landscape itself is never observed but only the dynamics of the experiment over the fitness landscape is observed. So, unlike the moment vectors for the rounds where we have sequencing data and we calculate $f^{(r)}$ as an estimate of $\mu^{(r)}$, we do not have an estimate for the moment vector of the fitness landscape $\mu$. To get around this, for each node $i$ or a pair of nodes $(i, j)$, we approximate the prevalence of codon $c$ at $i$ and the prevalence of the codon pair $(c,d)$ at $i, j$ at $r+1$ in terms of the moment vector $\mu^{(r)}$ of round $r$ and the moment vector of the fitness landscape $\mu$. In particular, assuming that the dependence between a set of nodes of interest $X_S$ and the remaining variables $X_{S^c}$ are limited for the approximation of $\mu_S^{(r+1)}$, we derive the following recursive relationships in (Sec. Approximate recursive relationships for marginals):

$$\begin{aligned} \mu_i^{(r+1)}(c) &\approx \sum_{c'\in\mathscr{C}} \frac{g_i(c'\to c)\mu_i(c)\mu_i^{(r)}(c')}{\sum_{c''\in\mathscr{C}} g_i(c'\to c'')\mu_i(c'')} \\ \mu_{ij}^{(r+1)}(c,d) &\approx \sum_{c',d'\in\mathscr{C}} \frac{g_i(c'\to c)g_j(d'\to d)\mu_{ij}(c,d)\mu_{ij}^{(r)}(c',d')}{\sum_{c'',d''\in\mathscr{C}} g_i(c'\to c'')g_j(d'\to d'')\mu_{ij}(c'',d'')}. \end{aligned} \tag{4}$$

We note that once $\mu_i^{(0)}$ is specified, all mean vectors $\mu_i^{(r)}$ are subsequently specified as functions of $\mu$ via the recursive relations in (4). Since at the start of the experiment, we have only copies of the wild-type sequence present (denoted by $w \in \Omega$) and so its frequency is 1 and the frequency of all other sequences is 0. Therefore, $\mu_i^{(0)}(c) = 1$ and $\mu_{ij}^{(0)}(c,d) = 1$ if $w_i = c$ and $w_j = d$ and it is 0 in all other cases.

Now, we estimate $\mu$ by minimizing the aggregated log-loss between the expected first and second-order frequencies $\mu_i^{(r)}$, $\mu_{ij}^{(r)}$ (as functions of $\mu$) and the observed first and second-order frequencies $f_i^{(r)}$, $f_{ij}^{(r)}$ for all positions $i, j \in [L]$, $i \neq j$. In doing so, we reparametrize $\mu$ so that we can optimize the objective function over amino-acid level parameters. We also enforce a local consistency condition, and regularize parameters to prevent overfitting during optimization (Sec. Inferring mean parameters). The canonical parameters of interest $\theta$ (which include the main effect parameters $\{h_i(a)\}$ and interaction (coupling) parameters $\{e_{ij}(a,b)\}$) are then estimated by regularizing and inverting the inferred covariance matrix from the estimated mean parameters. (Sec. Estimating canonical parameters).

We use the estimates of the canonical parameters of the Potts model (Main, Eq. 2) to get information about the protein's structure and function. Each parameter $e_{ij}(a,b)$ represents the interaction between amino acid $a$ at residue $i$ and amino acid $b$ at residue $j$. We compute an interaction score between residues $i$ and $j$ using the canonical parameters $e$ in the same manner as the DCA method [5]. The parameter set $e$ is over parameterized and so we first we convert to the zero-sum gauge and then we compute the Frobenius norm

$$e'_{ij}(a,b) = e_{ij}(a,b) - e_{ij}(\cdot,b) - e_{ij}(a,\cdot) + e_{ij}(\cdot,\cdot) \quad \text{and} \quad F_{ij} = \|e'_{ij}\|$$

The Frobenius norm $F_{ij}$ is the interaction score between pairs of residues $i$ and $j$. We only look at long range interactions (i.e. between residues that differ by more than 5 positions in the amino acid sequence).

### Approximate recursive relationships for marginals

Let $S \subseteq [L]$ be a set of nodes which is of interest for the approximation of $\mu_S^{(r)}$. We approximate $\pi$ with a distribution where the dependence between $X_S$ and the remaining variables $X_{S^c}$ is removed. In other words, we assume for any function $f$,

$$\mathbb{E}_\pi[f(X_{S^c})|X_S] \approx \mathbb{E}_\pi[f(X_{S^c})]. \tag{5}$$

Obviously, if $\pi$ factorizes, the approximation (5) is exact for any $S \subseteq [L]$. In general, the accuracy of approximation depends on the strength of interactions between $X_S$ and $X_{S^c}$.

$$\begin{aligned}
\mu_i^{(r+1)}(c) &= \sum_{x\in\Omega} p^{(r+1)}(x)\delta(x_i,c) \\
&= \sum_{x\in\Omega}\left\{\sum_{y\in\Omega} p^{(r)}(y)p(y\to x)\right\}\delta(x_i,c)(\text{by definition of } p^{(r+1)}(x)) \\
&= \sum_{x\in\Omega}\sum_{y\in\Omega} p^{(r)}(y)\cdot\frac{g(y\to x)\cdot\pi(x)}{\sum_{u\in\Omega} g(y\to u)\cdot\pi(u)}\cdot\delta(x_i,c)\,(\text{by definition of } p(x\to y)) \\
&= \sum_{y\in\Omega} p^{(r)}(y)\cdot\frac{\sum_{x\in\Omega} g(y\to x)\cdot\pi(x)\cdot\delta(x_i,c)}{\sum_{u\in\Omega} g(y\to u)\cdot\pi(u)}\ (\text{exchange of summation order}) \\
&= \sum_{y\in\Omega} p^{(r)}(y)\cdot\frac{g_i(y_i\to c)\,\mathbb{E}_\pi[\prod_{k\neq i} g_k(y_k\to X_k)\delta(X_i,c)]}{\sum_{c''\in\mathscr{C}} g_i(y_i\to c'')\,\mathbb{E}_\pi[\prod_{k\neq i} g_k(y_k\to U_k)\delta(U_i,c'')]}\ \text{where } X,U\sim\pi
\end{aligned} \tag{6}$$

Using the law of iterated expectations and $\mathbb{E}_\pi[\prod_{k\neq i} g_k(y_k\to X_k)|X_i] \approx \mathbb{E}_\pi[\prod_{k\neq i} g_k(y_k\to X_k)]$ by the assumption (5), we have,

$$\begin{aligned}
\mathbb{E}_\pi[\prod_{k\neq i} g_k(y_k\to X_k)\delta(X_i,c)] &= \mathbb{E}_\pi[\mathbb{E}_\pi[\prod_{k\neq i} g_k(y_k\to X_k)|X_i]\delta(X_i,c)] \\
&\approx \mathbb{E}_\pi[\prod_{k\neq i} g_k(y_k\to X_k)]\,\mathbb{E}_\pi[\delta(X_i,c)].
\end{aligned}$$

Therefore,

$$\mu_i^{(r+1)}(c) \approx \sum_{y\in\Omega} p^{(r)}(y)\cdot \frac{g_i(y_i \to c)\,\mathbb{E}_\pi[\prod_{k\neq i} g_k(y_k \to X_k)]\,\mathbb{E}_\pi[\delta\,(X_i,c)]}{\sum_{c''\in\mathscr{C}} g_i(y_i \to c'')\,\mathbb{E}_\pi[\prod_{k\neq i} g_k(y_k \to U_k)]\,\mathbb{E}_\pi[\delta(U_i,c'')]}$$
$$= \sum_{y\in\Omega} p^{(r)}(y)\cdot \frac{g_i(y_i\to c)\mu_i(c)}{\sum_{c''\in\mathscr{C}} g_i(y_i\to c'')\mu_i(c'')} \text{ since } \mu_i(c) = \mathbb{E}_\pi[\delta\,(V_i,c)] \text{ for } V\sim\pi$$
$$= \sum\nolimits_{c'\in\mathscr{C}} \frac{g_i(c'\to c)\mu_i(c)}{\sum_{c''} g_i(c'\to c'')\mu_i(c'')}\mu_i^{(r)}(c')$$

Note when $r=0$, $\mu_i^{(0)}(w_i)=1$ and $\mu_i^{(0)}(c')=0$ for $c'\neq w_i$. Therefore,

$$\mu_i^{(1)}(c) = \frac{g_i(w_i\to c)\mu_i(c)}{\sum_{c'} g_i(w_i\to c')\mu_i(c')}.$$

The derivation for the recursive relationships for pairwise marginals $\mu_{ij}^{(r)}$ are similar to the derivation above where we carry out similar calculations with $S=\{i,j\}$ instead of $S=\{i\}$, and is written in the next section.

#### Approximate recursive relationships for the second-order marginals

We have,

$$\mu_{ij}^{(r+1)}(c,d)$$
$$= \sum_{x\in\Omega} p^{(r+1)}(x)\delta\,(x_i,c)\,\delta\,(x_j,d)$$
$$= \sum_{x\in\Omega}\left\{\sum_{y\in\Omega} p^{(r)}(y)\cdot p(y\to x)\right\}\delta\,(x_i,c)\,\delta\,(x_j,d)\,\text{(by definition of } p^{(r+1)}(x))$$
$$= \sum_{x\in\Omega}\sum_{y\in\Omega} p^{(r)}(y)\cdot\frac{g(y\to x)\cdot\pi(x)}{\sum_{u\in\Omega} g(y\to u)\cdot\pi(u)}\cdot\delta\,(x_i,c)\,\delta\,(x_j,d)\,\text{(by definition of } p(x\to y))$$
$$= \sum_{y\in\Omega} p^{(r)}(y)\cdot\frac{\sum_{x\in\Omega} g(y\to x)\cdot\pi(x)\cdot\delta\,(x_i,c)\,\delta\,(x_j,d)}{\sum_{u\in\Omega} g(y\to u)\cdot\pi(u)}\text{ (exchange of summation order)}$$
$$= \sum_{y\in\Omega} p^{(r)}(y)\cdot\frac{\sum_{x\in\Omega} g(y\to x)\cdot\pi(x)\cdot\delta\,(x_i,c)\,\delta\,(x_j,d)}{\sum_{u\in\Omega}\sum_{c'',d''} g(y\to u)\cdot\pi(u)\cdot\delta\,(u_i,c'')\,\delta\,(u_j,d'')}$$
$$= \sum_{y\in\Omega} p^{(r)}(y)\cdot\frac{\mathbb{E}_\pi\left[\delta\,(X_i,c)\,\delta(X_j,d)\,g(y\to X)\right]}{\sum_{c'',d''}\mathbb{E}_\pi\left[\delta\,(U_i,c'')\,\delta\,(U_j,d'')\,g(y\to U)\right]}\text{ where } X,U\sim\pi$$
$$= \sum_{y\in\Omega} p^{(r)}(y)\cdot\frac{\mathbb{E}_\pi\left[\delta\,(X_i,c)\,\delta\,(X_j,d)\,g_i\,(y_i\to c)\,g_j\,(y_j\to d)\prod_{k\neq(i,j)} g_k(y_k\to X_k)\right]}{\sum_{c'',d''}\mathbb{E}_\pi\left[\delta\,(U_i,c'')\,\delta\,(U_j,d'')\,g_i\,(y_i\to c'')\,g_j\,(y_j\to d'')\prod_{k\neq(i,j)} g_k(y_k\to U_k)\right]} \tag{7}$$
$$\approx \sum_{y\in\Omega} p^{(r)}(y)\frac{g_i\,(y_i\to c)\,g_j\,(y_i\to d)\,\mathbb{E}_\pi\left[\delta\,(X_i,c)\,\delta\,(X_j,d)\right]\mathbb{E}_\pi[\prod_{k\neq(i,j)} g_k(y_k\to X_k)]}{\sum_{c'',d''} g_i\,(y_i\to c'')\,g_j\,(y_j\to d'')\,\mathbb{E}_\pi\left[\delta\,(U_i,c'')\,\delta\,(U_j,d'')\right]\mathbb{E}_\pi[\prod_{k\neq(i,j)} g_k(y_k\to U_k)]}. \tag{8}$$

where the equality in (7) is due to the definition $g(y \to x) = \prod_j g_j(y_j \to x_j)$, and the approximation in (8) is due to condition (5). Continuing to work with (8),

$$\begin{aligned}
(8) &= \sum_{y\in\Omega} p^{(r)}(y) \frac{g_i(y_i \to c)\, g_j(y_i \to d)\, \mathbb{E}_\pi\left[\delta(X_i,c)\,\delta(X_j,d)\right] \mathbb{E}_\pi[\prod_{k\neq(i,j)} g_k(y_k \to X_k)]}{\sum_{c'',d''} g_i(y_i \to c'')\, g_j(y_j \to d'')\, \mathbb{E}_\pi\left[\delta(U_i,c'')\,\delta(U_j,d'')\right] \mathbb{E}_\pi[\prod_{k\neq(i,j)} g_k(y_k \to U_k)]} \\
&= \sum_{y\in\Omega} p^{(r)}(y) \frac{g_i(y_i \to c)\, g_j(y_i \to d)\, \mu_{ij}(c,d)}{\sum_{c'',d''} g_i(y_i \to c'')\, g_j(y_j \to d'')\, \mu_{ij}(c'',d'')} \qquad (9) \\
&= \sum_{y\in\Omega} \sum_{c'} \sum_{d'} p^{(r)}(y) \delta(y_i,c')\, \delta(y_j,d') \frac{g_i(c' \to c)\, g_j(d' \to d)\, \mu_{ij}(c,d)}{\sum_{c'',d''} g_i(c' \to c'')\, g_j(d' \to d'')\, \mu_{ij}(c'',d'')} \\
&= \mathbb{E}^{(r)}\left[\sum_{c'}\sum_{d'} \delta(Y_i,c')\, \delta(Y_j,d') \frac{g_i(c' \to c)\, g_j(d' \to d)\, \mu_{ij}(c,d)}{\sum_{c'',d''} g_i(c' \to c'')\, g_j(d' \to d'')\, \mu_{ij}(c'',d'')}\right] \text{ (where } Y \sim P^{(r)}) \\
&= \sum_{c'}\sum_{d'} \mu_{ij}^{(r)}(c',d') \frac{g_i(c' \to c)\, g_j(d' \to d)\, \mu_{ij}(c,d)}{\sum_{c'',d''} g_i(c' \to c'')\, g_j(d' \to d'')\, \mu_{ij}(c'',d'')}
\end{aligned}$$

and the equality in (9) is due to $X \overset{d}{=} U$.

### Inferring mean parameters

Here, we describe the approximate moment-matching by minimizing the aggregated log-losses between the expected and observed first and second-order frequencies. In other words, we would like to solve the following objective function:

$$\underset{\mu\in\mathscr{M}\cap\mathscr{U}}{\arg\min} \left\{ -\sum_{r\in\mathscr{R}} \sum_{i\in[L]} \sum_{c\in\mathscr{C}} f_i^{(r)}(c) \log \mu_i^{(r)}(c;\mu) - \sum_{r\in\mathscr{R}} \sum_{\substack{i,j\in[L]\\ i\neq j}} \sum_{c,d\in\mathscr{C}} f_{ij}^{(r)}(c,d) \log \mu_{ij}^{(r)}(c,d;\mu) \right\}, \qquad (10)$$

where $\mathscr{M} := \{\mu \in \mathbb{R}^d; \exists\, p \text{ such that } \mathbb{E}_p[\phi(X)] = \mu\}$ and $\phi(x) := [\{\delta(x_i,c)\}_{i\in[L],c\in\mathscr{C}}, \{\delta(x_i,c)\delta(x_j,d)\}_{i,j\in[L],i<j,c,d\in\mathscr{C}}]$ is a sufficient statistic for $\pi_\theta$, and $\mathscr{U} := \{\mu \in \mathbb{R}^d; \mu_i(c) = \mu_i(c')\, \forall i \in [L], \forall c,c' \in \mathscr{C}$ such that $AC(c) = AC(c')$ and $\mu_{ij}(c,d) = \mu_{ij}(c',d')\, \forall i,j \in [L]\, \forall c,c',d,d' \in \mathscr{C}$ such that $AC(c) = AC(c')$ and $AC(d) = AC(d')\}$ In words, the set $\mathscr{M}$ corresponds to the set of the globally consistent mean vectors, i.e., all first-order and pairwise marginal probabilities that can be realized by some distribution over $\{0,1\}^d$ where $d$ is the dimension of the sufficient statistic $\phi$, and the set $\mathscr{U}$ corresponds to the set of mean vectors such that the mean values only depend on the amino acid values of an input sequence.

First, to optimize the objective over amino-acid level parameters, we reparameterize the mean parameters $\mu$ of the Potts model on $\Omega$ in terms of $\gamma := [\{\gamma_i(a)\}_{i\in[L],a\in\mathscr{A}}, \{\gamma_{ij}(a,a')\}_{i,j\in[L],i<j,a,a'\in\mathscr{A}}]$ as follows:

$$\begin{aligned}
\mu_i(c;\gamma) &= \frac{\nu_i(\mathrm{AC}(c);\gamma)}{\sum_{c'\in\mathscr{C}} \nu_i(\mathrm{AC}(c');\gamma)} \quad \text{and} \quad \nu_i(a;\gamma) = \frac{\exp\{\gamma_i(a))\}}{\sum_{a'\in\mathscr{A}} \exp\{\gamma_i(a')\}} \\
\mu_{ij}(c,d;\gamma) &= \frac{\nu_{ij}(\mathrm{AC}(c),\mathrm{AC}(d);\gamma)}{\sum_{c',d'\in\mathscr{C}} \nu_{ij}(\mathrm{AC}(c'),\mathrm{AC}(d');\gamma)} \quad \text{and} \quad \nu_{ij}(a,b;\gamma) = \frac{\exp\{\gamma_{ij}(a,b))\}}{\sum_{a',b'\in\mathscr{A}} \exp\{\gamma_{ij}(a',b')\}}.
\end{aligned} \qquad (11)$$

We also define $\nu_{ji}(b,a;\gamma) = \nu_{ij}(a,b;\gamma)$ for $j > i$.

Although the set $\mathscr{M}$ can be characterized by a finite number of linear inequalities, the number of linear inequalities grows fast depending on the dimension $d$, and in general, it is known to be extremely difficult to optimize even a linear objective over $\mathscr{M}$ unless the dimension $d$ is small [6]. We proceed by considering the relaxation of the optimization problem (10) by enforcing normalization conditions

$$\sum_{a\in\mathscr{A}} \nu_i(a) = 1,\ \sum_{a,b\in\mathscr{A}} \nu_{ij}(a,b) = 1,\ \forall i,j \in [L], i \neq j \qquad (12)$$

and local consistency conditions

$$\nu_i(a) = \sum_{b\in\mathscr{A}} \nu_{ij}(a,b), \forall a \in \mathscr{C},\ \forall i,j \in [L], i \neq j. \tag{13}$$

Note (12) is satisfied by the reparameterization (11). We add a following penalty term $\mathscr{P}(\gamma)$ with a langrange multiplier $\rho$ to promote the local consistency conditions (13)

$$\mathscr{P}(\nu) = \sum_{i=1}^{L} \sum_{a\in\mathscr{A}} \|\nu_i(a) - \frac{1}{L-1} \sum_{j\neq i} \sum_{b\in\mathscr{A}} \nu_{ij}(a,b)\|$$

Finally, to handle the high-dimensionality of $\gamma$, we add $\ell_2$-regularization terms $\mathscr{R}_{\text{main}}$ and $\mathscr{R}_{\text{int}}$ with hyper parameters $\lambda_{\text{main}}$ and $\lambda_{\text{int}}$:

$$\mathscr{R}_{\text{main}}(\gamma) = \sum_{i\in[L]} \|\gamma_i\|^2 \quad \text{and} \quad \mathscr{R}_{\text{int}}(\gamma) = 2 \sum_{i,j\in[L];i<j} \|\gamma_{ij}\|^2.$$

In summary, we solve the following optimization problem:

$$\begin{aligned}\widehat{\gamma} = \underset{\gamma_i(a);\gamma_{ij}(a,b)\in\mathbb{R}}{\arg\min} \{ &-\sum_{r\in\mathscr{R}} \sum_{i\in[L]} \sum_{c\in\mathscr{C}} f_i^{(r)}(c) \log \mu_i^{(r)}(c;\gamma) - \sum_{r\in\mathscr{R}} \sum_{\substack{i,j\in[L]\\ i\neq j}} \sum_{c,d\in\mathscr{C}} f_{ij}^{(r)}(c,d) \log \mu_{ij}^{(r)}(c,d;\gamma) \\ &+ \lambda_{\text{main}} \mathscr{R}_{\text{main}}(\gamma) + \lambda_{\text{int}} \mathscr{R}_{\text{int}}(\gamma) + \rho \mathscr{P}(\gamma)\}.\end{aligned} \tag{14}$$

We initialize the parameters $\gamma$ for optimization to be the log of the pairwise frequencies of the last round that we have sequencing data for. A small pseudocount is added to the frequencies so that this initialization procedure is defined for missing frequencies. We compute first order derivatives of the objective using automatic differentiation. Also, we use a gradient descent optimizer together with an early stopping rule.

After optimization, we get the individual and pairwise estimates $\widehat{\gamma}$ and can compute an estimate for the mean parameters $\widehat{\nu}, \widehat{\mu}$ using equation (11).

#### Estimating canonical parameters

Now, we describe how we convert the estimates for the mean parameters $\mu$ to the corresponding canonical parameters $\theta$. In other words, we would like to obtain $\widehat{\theta}$ in the set of parameters $\Theta$ such that $E_{\widehat{\theta}}[\phi] = \widehat{\mu}$.

First of all, it is a well known fact that such $\widehat{\theta}$ exists in $\Theta$ if $\widehat{\mu}$ is in the interior of the set of valid mean parameters $\mathscr{M}$, although in general this mapping is not available in closed forms. The important exception is Gaussian Graphical Model (GGM) [7, 8] where for a normal distributed $z \in \mathbb{R}^p$ with mean $m$ and variance $\Sigma$, i.e., $z \sim P_\theta$ such that $\theta = (\{h_i\}_{i\in[p]}, \{e_{ij}\}_{i,j\in[p],i\leq j})$ with $\mathbb{E}_\theta[z] = m, \mathrm{V}ar_\theta[z] = \Sigma$, we can explicitly derive the following mean parameters $(m,\Sigma)$ to canonical parameters $\theta = (\eta, J)$ relationships:

$$\begin{aligned} f_\theta(z) &\propto \exp\{-\frac{1}{2}(z-m)^\top \Sigma^\dagger (z-m)\} \\ &\propto \exp\{z^\top \eta - \frac{1}{2} z^\top J z\}. \end{aligned}$$

where $J = \Sigma^\dagger \in \mathbb{R}^{p\times p}$ is a pseudo-inverse matrix of $\Sigma$ and $\eta = \Sigma^\dagger m \in \mathbb{R}^p$. The use of the pseudo-inverse matrix is to handle the case where $\Sigma$ is not of full-rank. We have $\Sigma^{-1} = \Sigma^\dagger$ if rank($\Sigma$) = $p$.

In particular, when $z \in \{0,1\}^{Lq_a}$ is a one-hot encoding of a sequence, i.e., $z = [\{z_{(i,a)}\}_{i\in[L],a\in\mathscr{A}}]$ for $z_{(i,a)} = \delta(x_i, a)$, we have $z^2 = z$, and therefore,

$$\begin{aligned} f_\theta(z) &\propto \exp\{z^\top \eta - \frac{1}{2} z^\top J z\} \\ &= \exp\{\sum_{s\in[Lq_a]} (z_s \eta_s - \frac{1}{2} J_{ss} z_s^2) - \sum_{s,t\in[Lq_a],s<t} J_{st} z_s z_t\} \\ &= \exp\left\{ \sum_{i\in[L]} \sum_{a\in\mathscr{A}} z_{(i,a)} h_i(a) + \sum_{i\in[L]} \sum_{i<j} \sum_{a,b\in\mathscr{A}} z_{(i,a)} z_{(j,b)} e_{ij}(a,b) \right\} \end{aligned}$$

where we let $h_s := (\eta_s - \frac{1}{2}J_{ss})$, $e_{st} := -J_{st}$ for $s,t \in [Lq_a], s < t$, and $h_i(a)$ and $e_{ij}(a,b)$ refers to $(i,a)$th and $(i,a),(j,b)$th element of $h$ and $e$.

In the following, we will utilize the relationships $h_s = (\Sigma^{\dagger}m)_s - \frac{1}{2}\Sigma^{\dagger}_{ss}$ and $e_{st} = -\Sigma^{\dagger}_{st}$ to estimate corresponding canonical parameters from the estimated mean parameters. First of all, we compute the estimated covariance matrix for $\Sigma := \mathrm{Var}_{\theta}(z) \in \mathbb{R}^{Lq_a \times Lq_a}$ between the first order sufficient statistics based on the estimated $\widehat{\nu}$. We choose the states that are the least frequent for each position as reference states. For each $i, j \in [L]$ and $a, b \in \mathscr{A}$, we have,

$$\widehat{\Sigma}((i,a),(j,b)) = \begin{cases} \widehat{\nu}_{ij}(a,b) - \widehat{\nu}_i(a)\widehat{\nu}_j(b) & \text{if } i \neq j \\ \widehat{\nu}_i(a) - \widehat{\nu}_i(a)^2 & \text{if } i = j, a = b \\ -\widehat{\nu}_i(a)\widehat{\nu}_i(b) & \text{if } i = j, a \neq b \end{cases}$$

If the estimated mean vector $\widehat{\nu}$ is globally consistent, then the resulting $\widehat{\Sigma}$ would be semi-positive definite. However, since we solved a relaxed problem in (14), the estimated $\nu$ is not necessarily globally consistent. One consequence of this is that $\widehat{\Sigma}$ may have negative eigenvalues. We carry out a regularized inference to estimate $(h,e)$. Specifically, we let

$$\widehat{\Sigma}_{\lambda} = (\widehat{\Sigma} + \lambda \mathrm{I}), \tag{15}$$

and obtain $\{\widehat{h}_i(a)\}_{i\in[L],a\in\mathscr{A}}$ and $\{\widehat{e}_{ij}(a,b)\}_{i,j\in[L],i<j,a,b\in\mathscr{A}}$ using

$$\widehat{h} = \widehat{\Sigma}_{\lambda}^{-1}\widehat{\nu}_1 - \frac{1}{2}\mathrm{diag}(\widehat{\Sigma}_{\lambda}^{-1}) \quad \text{and} \quad \widehat{e} = -\widehat{\Sigma}_{\lambda}^{-1}, \tag{16}$$

where we define $\widehat{\nu}_1 := [\{\widehat{\nu}_i(a)\}_{i\in[L],a\in\mathscr{A}}]$ as the collection of first-order mean vectors at the amino-acid levels.

Finally, we estimate the Potts model energy function $\mathscr{E}(x)$ for a sequence $x = [x_1, \ldots, x_L]$ as follows

$$\widehat{\mathscr{E}}_{\lambda}(x) := -\left(\sum_{i\in[L]} \widehat{h}_i(AC(x_i)) + \sum_{i\in[L]}\sum_{i<j} \widehat{e}_{ij}(AC(x_i), AC(x_j))\right).$$

We use this energy function to design new proteins.We note the estimation procedure via regularizing diagonals (15),(16) is valid in the following sense: if $\|\widehat{\Sigma} - \Sigma\|_2 \to 0$, $\widehat{\mathscr{E}}_{\lambda}(x) \to \mathscr{E}(x)$ and $\widehat{e}_{ij}(a,b) \to e_{ij}(a,b)$ as $\lambda \to 0$, for any $x$ and $(i,j)$ such that $i < j$.
